# Plasticity of maternal environment dependent expression-QTLs of tomato seeds

**DOI:** 10.1101/2021.03.29.437558

**Authors:** Mark G. Sterken, Harm Nijveen, Martijn van Zanten, Jose M. Jiménez-Gómez, Nafiseh Geshnizjani, Leo A. J. Willems, Juriaan Rienstra, Henk W. M. Hilhorst, Wilco Ligterink, Basten L. Snoek

## Abstract

Seeds are essential for plant reproduction, survival, and dispersal. Germination ability and successful establishment of young seedlings strongly depends on seed quality and on environmental factors such as nutrient availability. In tomato (*Solanum lycopersicum*) and many other species, seed quality and seedling establishment characteristics are determined by genetic variation, as well as the maternal environment in which the seeds develop and mature. The genetic contribution to variation in seed and seedling quality traits and environmental responsiveness can be estimated at transcriptome level in the dry seed by mapping genomic loci that affect gene expression (expression QTLs) in contrasting maternal environments.

In this study, we applied RNA-sequencing to measure gene expression of seeds of a tomato RIL population derived from a cross between *S. lycopersicum* (cv. Moneymaker) and *S. pimpinellifolium* (G1.1554). The seeds matured on plants cultivated under different nutritional environments; i.e. on high phosphorus or low nitrogen. The obtained SNPs were subsequently used to construct a high-density genetic map. We show how the genetic landscape of plasticity in gene regulation in dry seeds is affected by the maternal nutrient environment. The combined information on natural genetic variation mediating (variation in) responsiveness to the environment may contribute to knowledge-based breeding programs aiming to develop crop cultivars that are resilient to stressful environments.

## Introduction

Seeds are essential for reproduction and dispersal of plants and function as survival structures to overcome harsh environmental conditions unfavourable for plant growth. Well-timed development and ripening of seeds, to ensure optimal seed performance and the ability to germinate in a permissive environment, are therefore essential for plant fitness. Successful germination strongly depends on seed performance, which is affected by environmental conditions, such as temperature, water availability, light conditions, and the nutrient status that the maternal plant experienced [1–5]. More specifically, seed performance/germination in species such as tomato and the model plant *Arabidopsis thaliana* is determined during seed development and maturation, and depends on temperature [6–8], photoperiod [9, 10], nutrient composition and levels [8, 11, 12]. Seed quality, germination and seedling establishment traits also have strong genetic determinants and (natural) genetic variation in quality traits, including Quantitative Trait Loci (QTLs), have been reported [8, 12–17].

Phosphate and nitrate are essential plant nutrients with profound effects on plant growth [18, 19] and seed performance/germination traits [8, 12, 15, 20]. In Arabidopsis it has been shown that seeds produced by plants fertilized with higher-than-normal levels of phosphate showed increased germination rates under stressful conditions [8]. Nitrate is known to have a strong effect on seed germination and seed dormancy in multiple plant species [21], with high concentrations of nitrate supplied to the mother plant leading to lower dormancy of the seeds [11]. This is attributed to nitrogen effects on the gibberellin/abscisic acid (GA/ABA) balance in the seeds; with higher endogenous nitrate levels resulting in lower ABA levels in seeds and hence, shallower dormancy [22]. In Arabidopsis, altered nitrate levels experienced by the mother plant also has a substantial effect on the levels of multiple metabolites and transcripts in the seeds, with a notable reduction in nitrogen metabolism-related metabolites and genes [23].

Tomato (*Solanum lycopersicum*) is one of the most important vegetable crops worldwide and is a model organism for research on fruit-bearing crops [24–27]. However, in the process of domestication, breeding selection and propagation, a substantial fraction of the genetic variation of the founder’s germplasms has been lost [26–28]. Moreover, due to a focus on fruit quality, resistance and yield traits, other desirable traits that have not been directly selected for have been lost over time in modern varieties. This includes several seed quality traits [28–32]. Trait variation loss can be restored by including wild cultivars/ancestors of modern commercial tomato such as *Solanum pimpinellifolium,* that represent a rich source of genetic variation, in breeding programs and in studies on tomato (quantitative) genetics [26, 33–37]. For instance, wild cultivars have been used in genetic screens and genome wide association studies (GWAS) to discover genomic loci and genes involved in variation in metabolic traits [38–42], insect resistance [43], floral meristem identity [31], trichome formation [44], and fruit shape and size [28, 34, 45]. In addition to GWAS, Recombinant Inbred Line (RIL) populations, derived from experimental crossing between *S. lycopersicum* and *S. pimpinellifolium,* are frequently used to uncover the effect of genetic variation on tomato traits [46–52], including various seed quality traits [12, 14, 15, 17, 53, 54].

The introduction and improved feasibility of diverse ∼omics techniques have accelerated studies into the molecular mechanisms underlying natural variation in tomato traits in the past two decades [55]. In particular, advances in transcriptomics techniques such as microarray analysis and later RNA-sequencing, have proven useful in this context, by enabling e.g. GWAS studies. Moreover, measuring gene expression in RILs has enabled expression-QTL (eQTL) analysis as a powerful tool to detect gene regulatory loci [56–61]. Combining the wealth of information obtained by mapping eQTLs enables (re)construction of regulatory networks underlying plant traits [56, 60, 62]. In addition, comparison of eQTL profiles from multiple environments may aid our understanding of how genetic variation shapes the effects the environment has on the appearance of phenotypes [57, 63, 64]. In plant (Arabidopsis) and worm (*Caenorhabditis elegans*) model systems it has been shown that especially *trans-*eQTLs are dynamic and can be highly specific for a certain environment [57, 63–68].

Although seed quality and seedling establishment characteristics are determined by both genetic variation and the maternal environment in which the seeds develop and mature [8, 12, 15], it is currently unknown if the maternal environment causes a perturbated eQTL landscape in the progeny seeds and how the nutrient environment of the mother plant affects these landscapes. We therefore followed an RNA-seq approach and quantified natural variation in mRNA levels in the dry seeds of a tomato RIL population from a cross derived from *S. lycopersicum* (cv. Moneymaker) and *S. pimpinellifolium* (G1.1554) parents [17, 49], that were cultivated either in a low nitrogen or a high phosphorus environment. In this work we first present a high-density RNA-seq-derived genetic map of tomato and subsequently we demonstrate how the genetic landscape of gene regulation of tomato dry seeds is affected by the nutritional environment of the mother plant.

Altogether, our detailed analysis of the genetic underpinning of plasticity in gene expression as responsiveness to the maternal environment, attributed to the progeny seeds, may contribute to knowledge-based breeding programs aiming to develop crop cultivars that are resilient to stressful environments, including production of high-quality seeds under sub-optimal environmental conditions.

## Results

### An RNA-seq-derived genetic map of tomato

We performed an RNA-sequencing experiment to uncover the interplay between genetic variation, the nutritional status of the maternal environment and mRNA abundances in progeny tomato seeds. The used seeds were derived from tomato RIL plants of a cross between *S. lycopersicum* (cv. Moneymaker; MM) and *S. pimpinellifolium* (G1.1554; PI) [17, 49] and their parental lines. All maternal plants were pre-cultivated on standard nutrient conditions and upon flowering transferred to either low nitrogen (LN) or high phosphate (HP) nutrition (∼100 RILs in total, ∼50 RILs in each environment) [15].

In addition to estimating expression differences among individuals, RNA-seq reads allowed for the identification of single nucleotide polymorphisms (SNPs) in transcribed genes of the parental lines and the RILs. These SNPs were subsequently used to construct high density genetic and physical maps of the RIL population, to facilitate QTL and eQTL mappings [13, 69]. In total, we detected 43,188 consistent SNPs between the parental lines. These SNPs were subsequently used to reconstruct the genotypes (*i.e.* determine the crossover locations) of the RILs in high detail (**Figure 1A**). Across our RIL set, a balanced distribution of the parental alleles was observed genome-wide, with the notable exception of chromosome 2, which had a substantial higher frequency of PI alleles (**Figure 1B**). Overall, 2,847 recombination (crossover) events were detected across the RIL population. As expected, the crossovers were found almost exclusively in euchromatic regions of the chromosomes, causing severe distortion between the physical and genetic maps, as described before [70](**Figure 1C**). On average, two recombination events were detected per RIL per chromosome. Altogether, the population size and recombination events provided 4,515 unique genetic markers and 4,568 distinguishable genomic loci/bins suitable for mapping, improving the previously available map [71] (**Supplementary table 1**). The detected loci had a size-range from 60 Mb to 1.7 Kb, with an average locus size of 180 Kb and a median of 11 Kb (**Supplementary table 2**). Given the high local recombination frequency, relatively small loci were overrepresented towards the chromosome tips (**Figure 1C**). Together, our dataset enables precise mapping of QTLs and eQTLs, especially towards the tips of the chromosomes.

**Figure 1:**
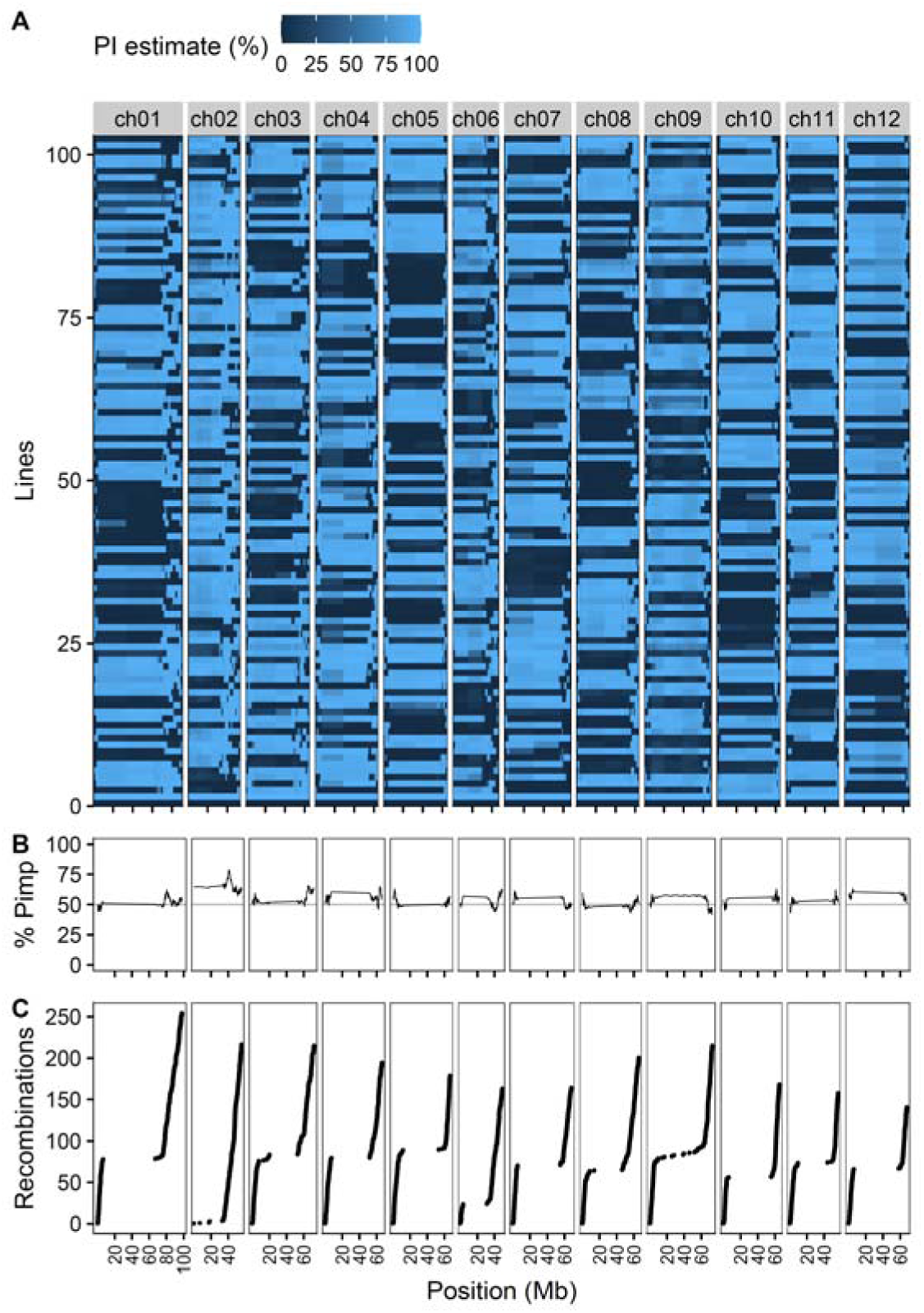
(**A**) Genetic map showing the genotype of the RILs and parental lines per chromosome (ch01 to ch12). Dark blue indicates MM (bottom horizontal line, with line number 0), light blue indicates PI (horizontal line above MM, with line number 100). Shades between dark and light blue visualize the certainty of the estimate that a locus corresponds to either MM or PI, depending on the SNPs identified (see legend above the panel; PI estimate). (**B**) Allele frequency (percentage) of *S. pimpinellifolium* (PI) alleles for each marker across the chromosomes, considering all RILs in the population. (**C**) Cumulative number of recombination events per chromosome for the whole population. Chromosome numbers are indicated above panels **A**, position on the chromosomes (in Mb) is shown on the x-axis below panel **C**.

**Table 1:**
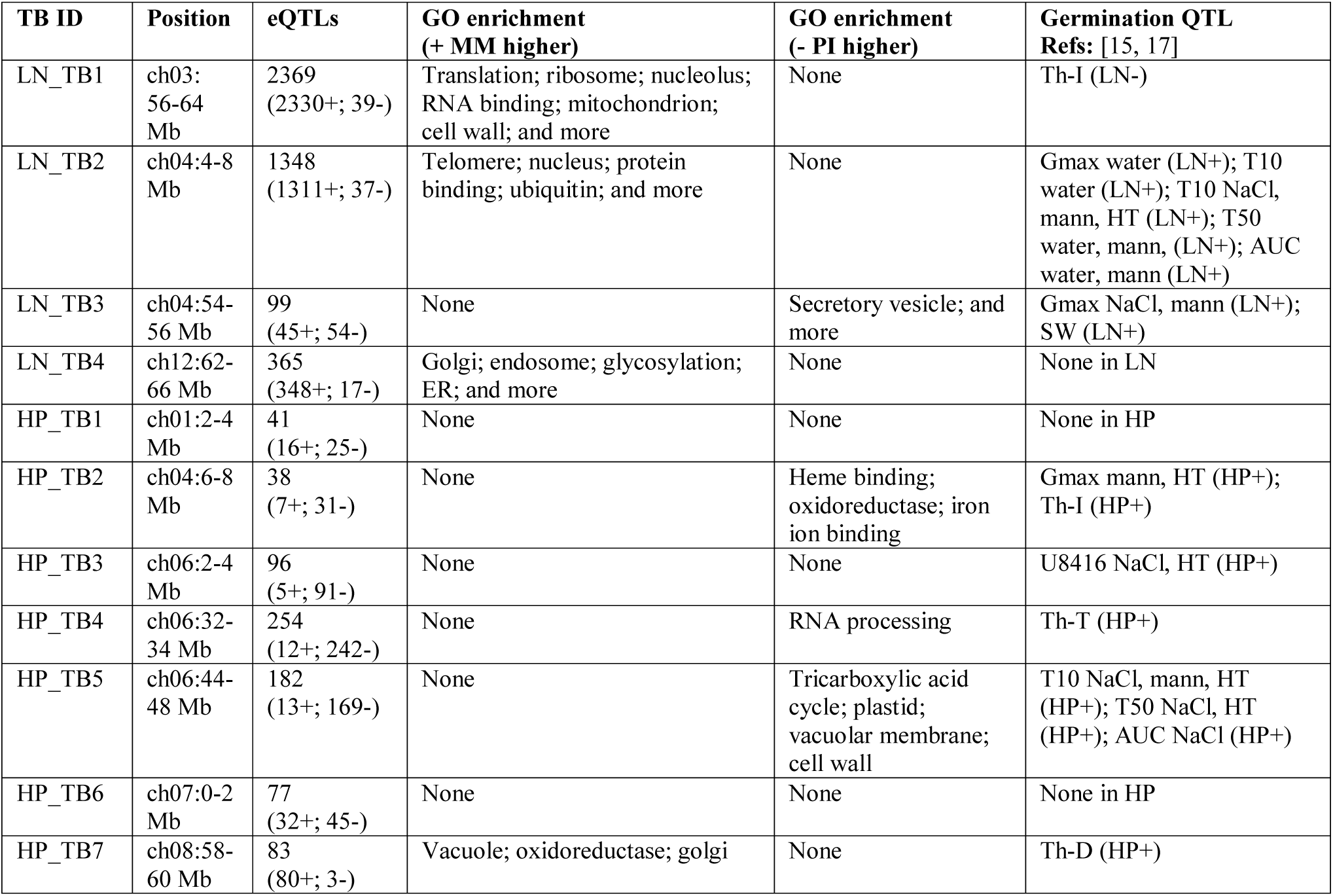
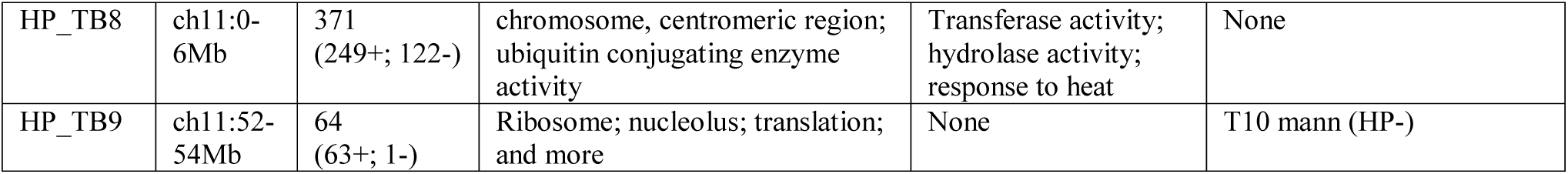
Overview of detected *Trans*-band (TB) eQTLs. Indicated are given ID’s, location on the physical genome (map position in Mb), number of eQTLs underlying the *trans*-band (+ sign: MM > PI; - sign PI > MM), GO terms enriched in the eQTLs underlying the *trans*-band in either MM or PI and co-location with known phenotypic QTLs for germination [15, 17]; + sign: MM > PI; - sign PI > MM).

### The maternal nutrient environment affects mRNA abundances in seeds

Next, we compared mRNA abundances in all HP-treated lines (RILs and parental lines) with the mRNA abundances in LN-treated lines, to identify genes contributing to differences between the two environments. Principal Component Analysis (PCA) demonstrated the presence of a substantial effect of the maternal nutrient environment on transcript levels in seeds (**Figure 2A**). A linear model was used to identify which mRNAs were differentially expressed between the two maternal environments. A multiple-testing correction was applied and differential expression of 2,871 mRNAs (out of 14,772 detected mRNAs) was found (Bonferroni corrected p-value < 0.05) to depend on the nutritional conditions the mother plant experienced during the seed maturation phase (i.e. LN or HP) (**Supplementary table 3**). Of these 2,871 mRNAs, 922 were more abundant in seeds developed and ripened in HP conditions compared to LN, and 1,949 mRNAs were significantly more abundant in LN conditions compared to HP. The mRNAs of genes that were more abundant after LN treatment were among others enriched for Gene Ontology (GO) terms: ‘chloroplast’, ‘ATP binding’, ‘proteasome’ and ‘nitrate transport’ (**Supplementary table 4**). mRNAs that were more abundant in seeds grown in HP conditions were enriched for the GO terms: ‘cellular response to hypoxia’, ‘pectin esterase activity’ and ‘glucosinolate metabolic process’ (**Supplementary table 4**).

**Figure 2:**
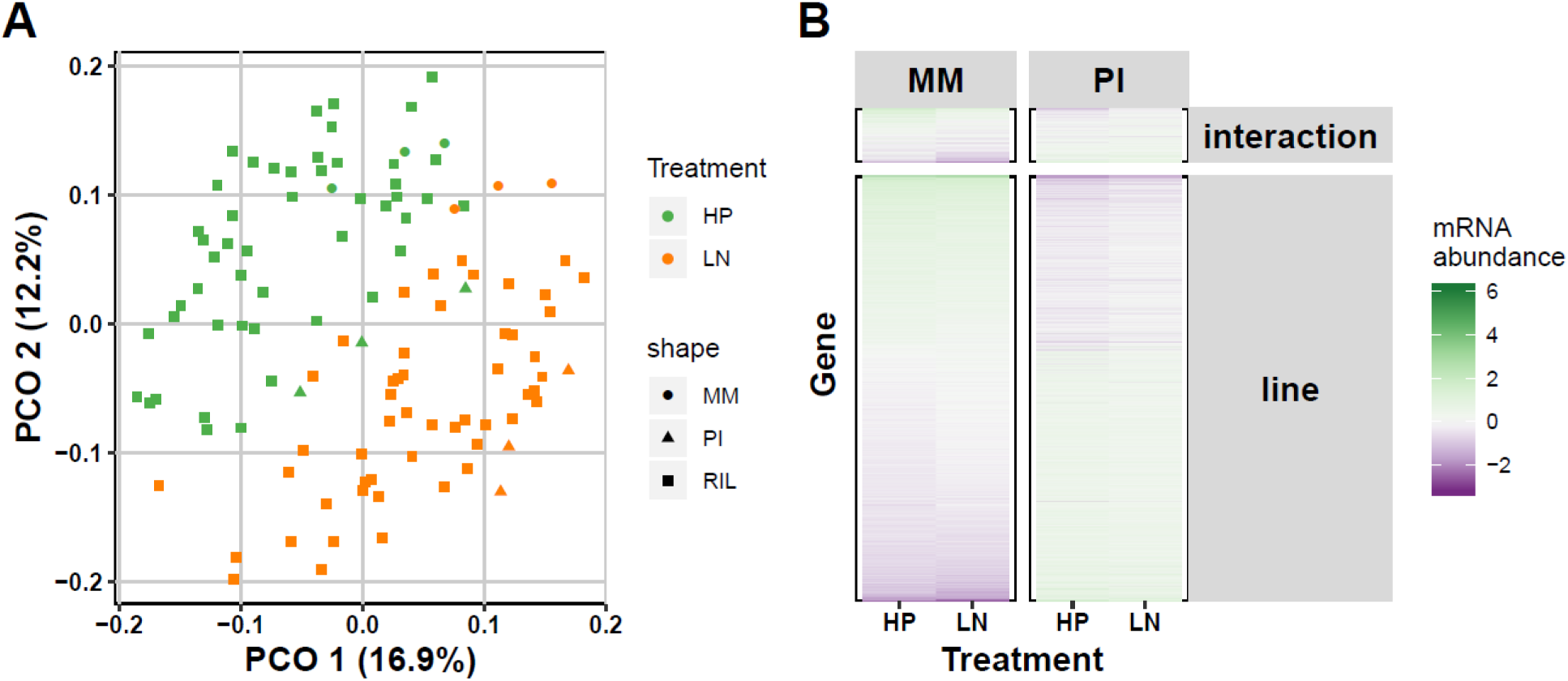
Nutrition status-related mRNA abundance differences between genotypes and seed development and maturation environments. (**A**) The first two axes of a principal component analysis on the log_2_ ratio with the mean transcripts per million (TPM) values. The first axis (PCO1) explained 16.9% of the variance in the data, the second 12.2%. Square symbols represent individual RILs, Moneymaker (MM) parental samples are represented by dots and *S. pimpinellifolium* (PI) parental samples by triangles. The colours indicate high phosphorous (HP; green) or low nitrogen (LN; orange) treatments applied to the mother plants. (**B**) Differentially abundant mRNAs in the two parental lines that are either not affected (line) or affected by treatment (interaction). Lower mRNA abundance is shown in purple and higher in green (see legend).

We also inquired the differences of the mRNA abundances between the MM and PI parental lines, within and between treatments. To this end, we again employed a linear model, but were less stringent in the statistical thresholds (as there were no confounding effects). We found 2,976 mRNAs differentially expressed between the two parental lines regardless of treatment and 382 mRNAs that were differentially expressed between the lines due to treatment (linear model, FDR ≤ 0.05; **Figure 2B** and **Supplementary table 5**). GO enrichment indicated that the 1,240 mRNAs more abundant in MM compared to PI were, among other categories, enriched, for ‘transcription factor activity’, ‘oxidation-reduction’, ‘protein-binding’, ‘-phosphorylation’, ‘-ubiquitination’, ‘chloroplast’, ‘circadian rhythm’, and ‘metal ion binding’ (**Supplementary table 6a**). The 1,736 mRNAs that were more abundant in PI compared to MM were, among other categories, enriched for ‘cytosol’, ‘chloroplast’, ‘nucleus’, ‘mitochondrion’, ‘cytoplasm’, ‘ribosome’, ‘translation’, ‘nucleolus’, ‘endoplasmic reticulum’, ‘oxidation-reduction’, ‘vacuole’, and ‘copper ion binding’ (**Supplementary table 6a**). The 382 genes showing a significant interaction effect between the parental background and maternal environment showed an enrichment for the GO terms ‘oxidation-reduction’, ‘extracellular region’, ‘transcript regulation’, ‘iron ion binding’, and ‘response to gibberellin’ (**Supplementary table 6b**). Of note, the ‘oxidation-reduction process’ and ‘transcript regulation’ GO terms are enriched in the upregulated genes of both MM and PI, which is not surprising since both GO terms are quite general and each represents many genes. These results show that the nutrition status of the mother plant (environment; E) as well as genotype (G), and the interaction between the two (G x E), modulate mRNA abundances in dry seeds of tomato.

### Heritability and transgression in mRNA abundances

To estimate the contribution of genetic variation to differences in mRNA abundance between the genetic backgrounds (plant lines) and treatments (nutrient status), we calculated the Broad-Sense Heritability (BSH). In addition, replicated measurements in the parental lines were used to estimate non-genetic variance. We found 5,112 genes in HP and 5,332 genes in LN that showed significant heritability for mRNA abundance, of which 2,973 genes overlapped (39.8%; permutation, FDR < 0.05; **Figure 3A; Supplementary table 7a**). Subsequently, we checked if genes with significant heritable contribution to mRNA abundance differences were predominantly affected by the maternal nutrient environment. However, we did not find such an enrichment for any of the overlapping groups of genes (hypergeometric test, p > 0.01; **Supplementary** **figure 1A**). We thus conclude that, overall, the number of genes with significant heritability for mRNA abundance were not specifically responsive to the maternal nutrient treatments. The genes with heritable mRNA abundance in HP alone were enriched for the GO terms: ‘translation’, ‘ribosome’, ‘mitochondrion’, and more (**Supplementary table 7b**). Those that showed significant heritability only in LN were enriched for the GO terms: ‘ABA metabolic process’, and others (**Supplementary table 7b**). The genes that showed significant heritability in both environments were enriched for various GO terms: ‘oxidation-reduction process’, ‘ribosome/translation’, ‘nucleolus’, ‘cell wall’, ‘heme binding’, ‘ion binding’, and ‘vacuole’ (**Supplementary table 7b**).

**Figure 3:**
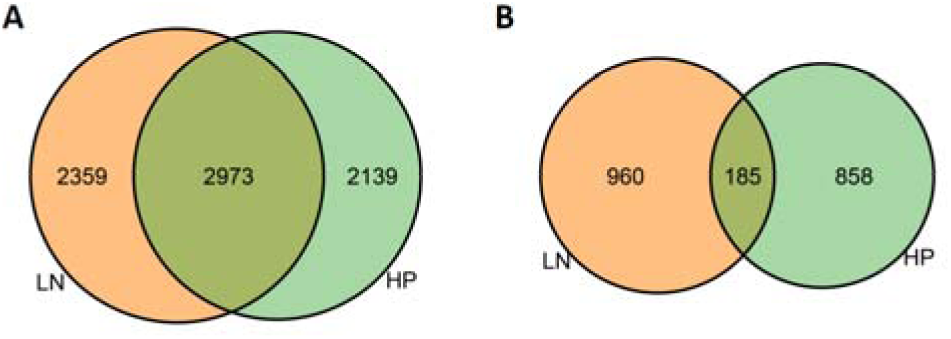
Venn diagrams showing the overlap and differences of **(A)** genes with significant heritable variance (Supplementary table 7a) and **(B)** genes exhibiting significant transgression (Supplementary table 8a), of mRNA abundance levels between LN (orange) and HP (green; FDR < 0.05).

We next assessed the complexity of the genetic regulation underlying mRNA abundance differences. To this end, the transgression was calculated, i.e., trait values in RILs that extent beyond the parental means. We found significant transgression in mRNA abundance (trait) levels for 1,043 genes in the maternal HP treatment and 1,145 genes in the maternal LN treatment (permutation, FDR < 0.05; **Supplementary table 8a**). This suggests a polygenic genetic architecture for mRNA abundance. Of these, the mRNA abundances of 185 genes showed significant transgression beyond the parental means in both treatments (**Figure 3B**). Also, here, we tested for significant overlap with treatment-related genes. Yet, with 18% response to treatment of the transgressive mRNAs, there was no significant enrichment for transgressive mRNA abundances with treatment-related differences (hypergeometric test, p > 0.01; **Supplementary** **figure 1B**). So, alike heritability, transgression is apparently not linked to a reduction of nitrogen or increase of phosphorus content in the maternal growth environment. Moreover, compared to genes showing significant heritability, many fewer GO terms were enriched in the genes showing transgression, and those GO terms that were enriched, generally had a lower level of significance. For genes showing transgression in HP alone, the GO terms ‘cell periphery’, ‘positive gravitropism’, ‘cysteine biosynthetic process’, ‘symporter activity’ and ‘response to heat’ were enriched. Whereas for genes only showing transgression in LN the GO-terms ‘beta-glucosidase activity’, ‘preprophase band’, and ‘phragmoplast’ were enriched. The GO term ‘DNA-binding transcription factor activity’ was enriched in genes showing transgression in both environments (**Supplementary table 8b**).

### The maternal nutrient environment produces specific eQTL landscapes

Altogether, our analyses revealed both a considerable effect of the maternal nutrient environment (HP versus LN) and a significant influence of genetic variation in the RIL panel (heritability) on the detected mRNA abundance levels. By combining our constructed high density SNP genetic map (**Figure 1A, Supplementary table 1**) with the obtained mRNA abundance dataset (**Figure 2**), we were able to identify eQTLs that potentially contribute to the variation in mRNA abundance (**Figure 4A-F)**. In other words, the identified eQTL loci have a high chance of harboring polymorphic regulatory factors (e.g., genes or other genetic elements) for mRNA abundance, prospectively explaining variation in the seed and germination trait phenotypes observed.

**Figure 4:**
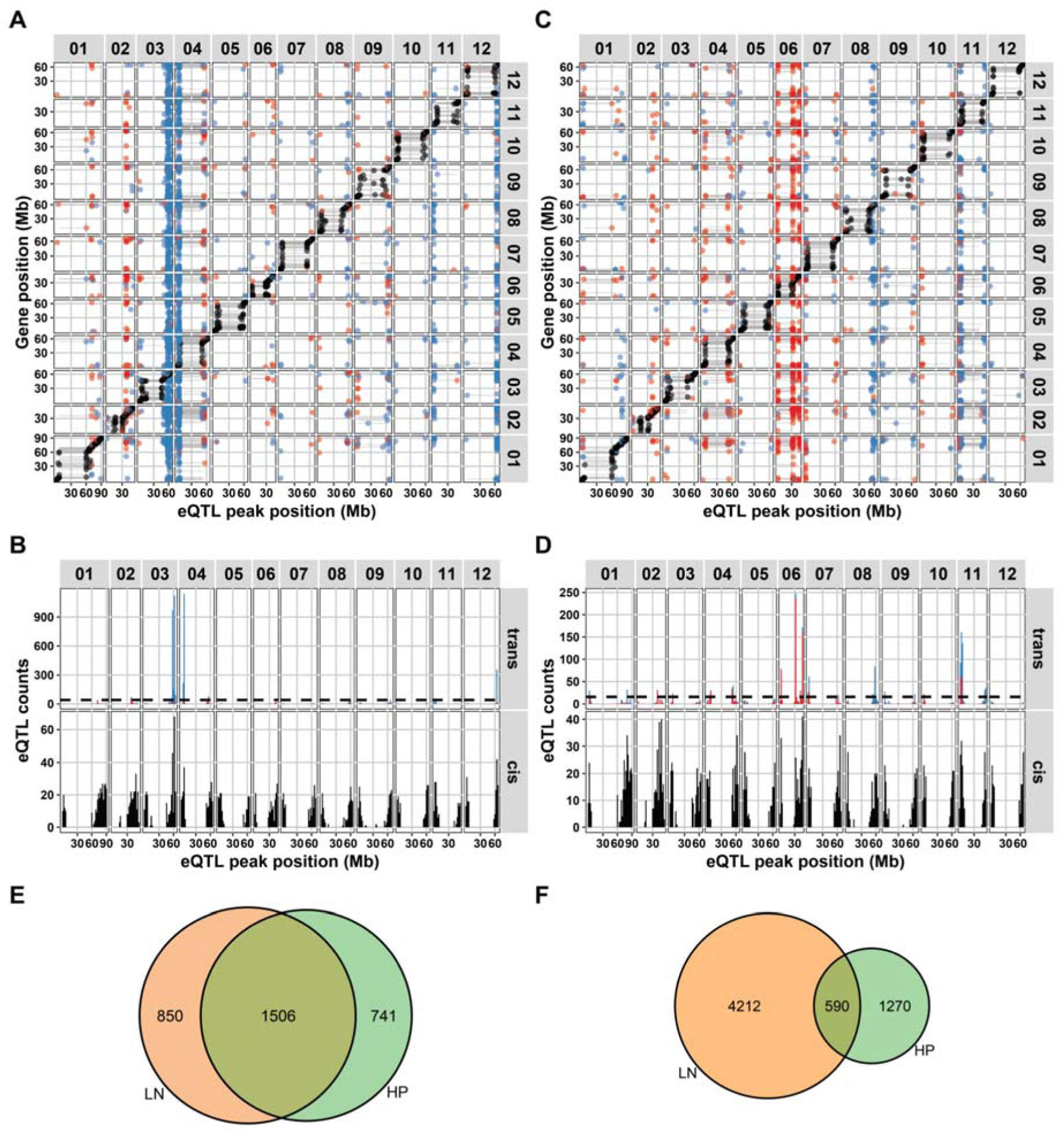
Characteristics of the detected eQTL landscapes in tomato dry seeds in (**A, B**) LN and (**C, D)** HP environments. (**A, C**) *Cis-trans* plots of eQTLs mapped (-log_10_(p) > 3.9). The positions (in Mb, per chromosome) of the eQTL peaks are plotted on the x-axis and the positions of the corresponding genes on the y-axis. Chromosome numbers are indicated on the top and right (grey labels). Colored dots indicate *cis*-eQTLs (black), eQTLs associated with higher mRNA abundance due to the MM allele (blue dots) or with higher abundance by the PI allele (red dots). (**B, D**) Histograms showing the distribution of the amount of *cis*-(lower panel) and *trans* (upper panel)-eQTLs over the chromosomes, arranged by eQTL peak location counted per 2 million bases (Mb) bins. The dashed lines in the *trans*-eQTL panels indicate the threshold for calling a *trans*-band (poisson distribution, p < 0.0001). (**E**) The overlap of *cis*-eQTLs in the two treatments and (**F**) the overlap of *trans*-eQTLs in the two maternal environments.

We detected a maternal environment-specific *trans*-eQTL landscape, as the distribution of the position of the *trans*-eQTLs was very different between the two environments. For the HP environment, 4,281 eQTLs for 3,833 genes were identified, of which 2,247 were *cis*-eQTLs and 2,034 were *trans*-eQTLs. For the LN environment, 7,487 eQTLs were detected for 6,815 genes, of which 2,356 were *cis*-eQTLs and 5,131 were *trans*-eQTLs (FDR < 0.05; −log_10_(p) > 3.9; **Figure 4A-D; Supplementary Table 9**; **Supplementary Table 10**). A significant overlap between *cis*-eQTLs of the two environments was noted (**Figure 4E;** 1,506 overlapping *cis*-eQTLs; 48.6%; hypergeometric test, p < 1*10^-16^). On the contrary, the *trans*-eQTLs were mainly specific for each tested maternal environment (**Figure 4F**; 590 overlapping *trans*-eQTLs; 9.7%; hypergeometric test, p = 1.0). However, both *cis-* and *trans*-eQTLs were not enriched for genes with differentially abundant mRNA levels based on the maternal environment (hypergeometric test, p > 0.01; **Supplementary** **figure 2A** and **B**). Together with the significant transgression (**Supplementary table 8a)** and considerable heritability of mRNA abundances (**Supplementary table 7a; Figure 3A)**, this indicates that *trans*-eQTLs represent a genotype-specific interaction with the maternal nutrient environment. Many different GO terms were found to be enriched in the genes with environment specific eQTLs. For an overview see **Supplementary table 11**).

The majority of the *trans*-eQTLs clustered in maternal nutrient environment-specific eQTL hotspots or *trans*-bands (**Figure 4A, C**). Hence, these genomic regions harbor the main loci underlying the genetic variation in environment-specific gene expression regulation in our dataset. A total of 13 *trans*-bands (9 in the HP treatment and 4 in the LN treatment; see Methods for the *trans*-band criteria) were identified, which account for 1,206 of the *trans-*eQTLs in the HP treatment (59.3% of HP total) and 4,181 of the *trans*-eQTLs in the LN treatment (81.5% of LN total; **Table 1**).

Thus, *trans*-bands are a major explanatory factor for *trans*-eQTLs. In other words, a relatively large proportion of *trans*-eQTLs are caused by a few pleiotropic major-effect loci. Remarkably, the MM allele had a positive effect on mRNA abundance for the majority of the eQTLs of the *trans*-bands in the LN soil environment, whereas this was not so prevalent in the HP environment (**Table 1; Figure 4A, C**). Most of these *trans*-bands showed enrichment for specific GO terms, such as ‘translation’ and ‘specific cellular organelles’ for LN and, ‘oxidoreductase’ and ‘vacuole’ for HP. (**Table 1**, **Supplementary table 12**). Moreover, many of the *trans*-bands co-locate with known QTLs for germination and seed traits (**Table 1** [15, 17]). These eQTLs can therefore contribute to uncovering the molecular genetic mechanisms underlying the germination and seed trait QTLs.

## Discussion

Our RNA-sequencing data obtained from a Tomato RIL population (*S. lycopersicum* (cv. Moneymaker; MM) x *S. pimpinellifolium* (G1.1554; PI)) [17, 49], allowed for the construction of a detailed and high resolution genetic map, describing the genotypes using 4,515 SNP markers. This is over five times more than previously reported in Kazmi *et al*., 2012 [71], which used 865 markers. However, intrinsic to RNA-seq data, only SNPs present in the coding parts of the genes (mRNA’s) could be used. Therefore, determining the exact locus where recombination took place would need additional genome sequencing as described in [70].

By measuring transcript levels (i.e. mRNA abundances) in the seeds of a tomato RIL population that had matured in different maternal nutrient environments, we show that the maternal environment affects both regulation and the genetic architecture of gene expression in progeny seeds. Especially, *trans* eQTLs proved environment specific, which is comparable to other species [57, 63–67, 72–74]. We found 3,833 genes (∼26% of all detected expressed genes in the RILs), with an eQTL in HP and 6,815 genes (∼46% of all expressed genes in the RILs) with an eQTL in LN. This is comparable to the number detected by Ranjan *et al*. 2016 [75], who used the upper part of 5 day-old hypocotyls of introgression lines (ILs), developed from the wild desert-adapted species *Solanum pennellii* and domesticated *Solanum lycopersicum* cv. M82 [76], and found 5,300 genes (∼25% of total expressed genes) to have an eQTL, with roughly half in *cis* and half in *trans*. We also found this close to 50/50 ratio in the HP condition, whereas in the LN condition the ratio of *cis/trans* eQTLs was increased to 30/70. Research in yeast indicated that the detection of *trans*-acting eQTLs is more strongly affected by the power of the study than detection of *cis*-acting eQTLs [73]. So, it is likely that in our study we would have even more *trans*-eQTLs relative to *cis-*eQTLs.

By comparing two different maternal environments in a population originating from two different genetic backgrounds, many different maternal environment specific eQTLs were detected. This underlines the interplay between genetics and nutrient environment in our study. Yet, we expect much of the variation caused by this interplay will be uncovered in future studies increasing numbers of different timepoints, environments and genotypes. More detailed data on the number and type of polymorphisms between tomato lines, such as frameshifts [77] and copy number variations [35], could facilitate identification of the causal polymorphic genes in this and other eQTL studies. Moreover, combining eQTLs with QTLs obtained using phenotypic trait data [12, 15, 17], as well as other molecular data such as proteomics and/or metabolomics [47], will contribute to obtaining mechanistic insight on how genotypic variation leads to phenotypic variation between individuals at a systemic level. Furthermore, these eQTLs could be used as a lead in studies with a larger source of wild-genotypes and combined with GWAS [39-42, 44, 78], to pinpoint causal polymorphisms underlying variation at both the molecular and phenotypic levels.

## Methods

### Plant lines, growth conditions, and nutrient treatments

The mother plants (maternal conditions) were cultivated as described in Kazmi *et al*. 2012 and Geshnizjani *et al*. 2020 [15, 17], in the greenhouse at Wageningen University, the Netherlands. In short; the parental lines *Solanum lycopersicum* cv. Money maker (MM) and *Solanum pimpinellifolium* accession CGN14498 (PI) as well as the derived recombinant inbred lines (RILs; [49]; **Supplementary table 1**) were grown on rockwool under standard nutrient conditions (14 mM Nitrate and 1 mM Phosphate) with a 16h light (25°C) and 8h darkness (15°C) photoperiod. From the moment the first flower opened, the plants were fertilized with the specific nutrient solutions, low nitrate (2.4 mM Nitrate, 1 mM Phosphate) and high phosphate (14 mM Nitrate, 5 mM Phosphate) in two biological replicates per environment. The seeds were collected from healthy and ripe fruits and the pulp still attached to the seeds was removed with 1% hydrochloric acid (HCl) and a mesh sieve. Water was used to remove the remaining HCl and pulp. For disinfection, seeds were treated with trisodium phosphate (Na3PO4.12H2O). Subsequently, seeds were dried at 20°C for 3 days on a clean filter paper in ambient conditions. The seeds were then stored in paper bags at room temperature.

### RNA-isolation, library prep and RNA-seq

We used 10 mg grinded powder derived from 30 whole, dry, brushed, after-ripened seeds (12 months after harvest) of parental lines and the RILs grown under the different nutrient environments in a GGG design [79, 80] to extract total RNA. In total, 3 replicates per treatment for the parental lines where sequenced and 49 single RIL seed pools for HP and 52 single RIL seed pools for LN (**Supplementary table 13**). RNA was isolated using the NucleoSpin RNA plant isolation kit (Macherey-Nagel 740949) with on-column DNA digestion and adding Plant RNA isolation Aid (Life technologies) according to the manufacturer’s protocol and instructions. Strand-specific RNA-seq libraries were prepared from each RNA sample using the TruSeq RNA kit from Illumina according to manufacturer’s instructions. Poly-A-selected mRNA was sequenced using the Illumina HiSeq2500 sequencer, producing strand-specific single-end reads of 100 nucleotides. Raw sequence reads can be found in the Sequence Read Archive (SRA; www.ncbi.nlm.nih.gov/sra) under ID: PRJNA704909

### Alignment and SNP calling

Reads were trimmed using Trimmomatic (version 0.33, [81] to remove low quality nucleotides. Trimmed reads were subsequently mapped to the Tomato SL4.0 reference genome with the ITAG4.0 annotation [82] using the HISAT2 software (version 2.1.0, [83] with the --dta-cufflinks option. The resulting SAM alignment files were sorted and indexed using samtools version1.9 [84]. SNPs were called using bcftools mpileup with a minimum read depth of 3.

### Generation of a genetic map from RNA-seq data

The genetic map used for mapping the eQTLs was made from the RNA-seq data following the protocol described in Serin & Snoek *et al*. 2017 [13] and Snoek *et al.* 2019 [69]. With the following modifications: SNPs were filtered for those that were consistently found in all replicates of the parental lines and observed in all RILs. Then the genotype per RIL was determined per sliding bin of 100 SNPs where the mean position of those SNPs was taken as the physical position of the obtained marker.

### Quantification of RNAseq

Before mRNA abundance analysis, between 12M and 31M reads per sample were mapped to the SL4.0 genome with ITAG4.0 annotation [82] using HISAT2 as described above. The mRNA abundance was quantified to counts using Stringtie [85] with the options -e, -B and -G. In R, the counts were used to calculate transcripts per million (TPM). The TPM values were log_2_-transformed by

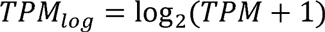

Additionally, to use for statistics, also a ratio with the average was calculated, by

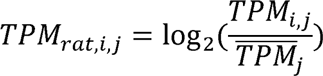

Where the log_2_ was calculated for each transcript *i* of sample *j* by dividing over the average value for that transcript 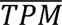 over all samples *j*. After transformation, the transcripts were filtered for TPM_log_ > 0, and detection in all samples.

### mRNA abundance analysis and QTL analyses

The analyses reported below were conducted in “R” (version 3.5.3, x64)[86] with custom written scripts, accessible via https://git.wur.nl/published_papers/sterken_tomato-eqtl_2021. For analysis, the dplyr and tidyr packages were used for data organization [87, 88], and plots were generated using ggplot2 [89].

### Treatment related mRNA abundance differences

The principal component analysis comparing the mRNA abundances was done on the *TPM_rat_*-transformed data, using the *prcomp* function in “R”. The mRNA abundance differences between treatments were tested between the LN and HP treatments using the linear model

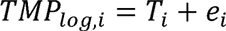

where *TPM_log,i_* is the abundance level of transcript *i* (one of 14,772 transcripts) in RIL *j* (n = 55 for the HP treatment and n = 58 for the LN treatment), *T* is the treatment (HP or LN), and *e* is the error term. To reduce the chance of detecting differences due to genetic variation, a strict multiple-testing correction was applied (Bonferroni) using *p.adjust*. The threshold for significance was −log_10_(p) > 5.47 (FDR = 0.05).

To determine the effect of treatment on the differences in mRNA abundance between the parental lines, we ran a linear model explaining the differences due to treatment and line effects on the MM and PI parental data. The model used was

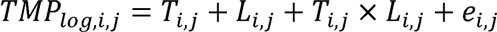

where *TPM_log,i,j_* is the abundance level of transcript *i* (one of 14,772 transcripts) in parental replicate *j* (n = 3 for both treatments for MM and PI), *T* is the treatment (HP or LN), *L* is the line (MM or PI), and *e* is the error term. Values were corrected for multiple testing using *p.adjust* following the Benjamini Hochberg algorithm. The thresholds for FDR = 0.05 were: −log_10_(p) = 1.71 for line, −log_10_(p) = 2.08 for treatment, and −log_10_(p) = 2.89 for the interaction between line and treatment. We took the most stringent p value, −log_10_(p) = 2.89 as threshold to determine significance.

### Transgression

Transgression was calculated by counting the number of lines with expression levels beyond three standard deviations from the mean of the parental lines (as in RB Brem and L Kruglyak [90]); µ ± 3*σ. This was done for both treatments separately. The lower boundary was established by the parental line with the lowest mean, and the upper boundary was established by the parental line with the highest mean. The standard deviation used to determine transgression (σ) was calculated as the pooled standard deviation of the two parental lines (n =3 for both).

Significance of the transgression was calculated by permutation. The expression values were randomized over the line designations and the same test as above was conducted. This was repeated 1000 times for each transcript, so the obtained values could be used as the by-chance distribution. The 50^th^ highest value was used as the false discovery rate (FDR) = 0.05 threshold.

### Heritability

The heritability was calculated by estimating the genotypic variance in the RILs and the remaining variance (e.g. measurement error) in the parental lines (as in JJ Keurentjes, J Fu, IR Terpstra, JM Garcia, G van den Ackerveken, LB Snoek, AJ Peeters, D Vreugdenhil, M Koornneef and RC Jansen [56]). This was done for both treatments separately, by

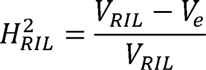

where *V_RIL_* is the variance within the RIL population and *V_e_* is the pooled variance of both parental lines.

To establish whether the heritability was significant and not outlier-driven, we applied a permutation approach (as in A Vinuela, LB Snoek, JA Riksen and JE Kammenga [91]). The trait values were randomized over the line designations and the heritability calculation were repeated. This was done 1000 times for each transcript to generate a by-chance distribution. The 50^th^ highest value was used as the FDR = 0.05 threshold.

### eQTL mapping

For eQTL mapping a single marker model was used, and was applied separately for both treatments (as in [65, 92]). QTLs were mapped using the model

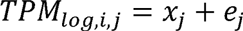

where *TPM_log,i,j_* is the expression level of transcript *i* (one of 14,772 transcripts) in RIL *j* (n = 49 for the HP treatment and n = 52 for the LN treatment). The expression levels were explained over the genotype on marker location *x* (x = 1, 2, …, 4515) of RIL *j*.

To determine the reliability of the detected QTLs and correct for multiple testing, a permutation approach was used. As in the other permutations, the expression levels were randomly distributed over the lines and this randomized set was mapped again according to the procedure described above, which was repeated 10 times. To determine the FDR, we applied a correction for multiple testing under dependency [93]

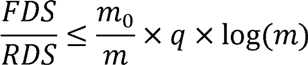

where *FDS* (false discovery) is the number of eQTLs detected in the permutation and the *RDS* (real discovery) is the number of eQTLs detected in the QTL mapping at a specific significance level. The number of true null hypotheses tested (*m_0_*), was 14,772 -*RDS*, where the number of hypotheses tested (*m*) was the number of transcripts, 14,772. The q-value was set at 0.05, which led to a threshold of –log_10_(p) = 3.7 for the LN treatment and –log_10_(p) = 3.9 for the HP treatment. To keep comparisons straightforward (similar effect sizes), analyses were conducted at the most stringent threshold (-log_10_(p) > 3.9).

The eQTL types (*cis* or *trans*) were called based on distance to the gene encoding the affected transcript. A *trans*-eQTL had to be located at least 1 Mb from the gene. Furthermore, we calculated the confidence interval of the QTL as a 1.5-drop from the highest –log_10_(p). For a *trans*-eQTL to be called, the location of the affect transcript was required to be outside of this confidence interval as well.

### Trans-band identification

Identification of regulatory hotspots (*trans*-bands) was based on assessing whether the number of *trans*-eQTLs mapped to a locus exceeded the expected number based on an equal genome-wide distribution (as in [65, 94]. We used a Poisson distribution to ascertain the significance of eQTL abundances per 2 Mb bin. For the HP treatment, we expected 15.8 *trans*-eQTL per bin, and for the LN treatment we expected 40.8 *trans*-eQTL per bin. We used a conservative threshold for calling a bin enriched in *trans*-eQTL, p < 0.0001. After identifying significant bins, adjacent bins (significant bins, with up to 1 non-significant bin in-between) were merged to a single *trans*-band.

### Enrichment

GO enrichment was determined using the hypergeometric test in R on the GO annotation done for ITAG2.4 downloaded from AgriGO (www.bioinfo.cau.edu.cn/agriGO) [95] combined with the annotation for ITAG3.1 and expanded with the GO annotation of the Arabidopsis homologues. All expressed genes were used as background genes in the enrichment test.

### Map and eQTL data in TomQTL

The physical map of the RIL population and the eQTL –log10(p-value) scores are available for download and online exploration in TomQTL at http://www.bioinformatics.nl/TomQTL/, an interactive website based on AraQTL [63] and WormQTL2 [74].

## Acknowledgements

We thank Prof. Dr. Jan Kammenga of the Laboratory of Nematology of Wageningen University, Prof. Dr. Dick de Ridder of the Laboratory of Bioinformatics of Wageningen University and Prof. Dr. Berend Snel of the Theoretical Biology and Bioinformatics department of Utrecht University for their support.

## Author contributions

WL and HWMH conceived the study, NG, JR and LW performed the experiments, MGS, HN and LBS analyzed and visualized the data, MGS and LBS wrote the manuscript with input from MvZ, HN, JJG, and WL. All authors approved the final version of this manuscript.

## Funding

This work was supported by Technology Foundation (STW), which is part of the Netherlands Organization for Scientific Research (NWO) (LW, JR, HN, WL). M.G.S. was supported by NWO domain Applied and Engineering Sciences VENI grant (17282).

## Supplementary tables and figures

**Supplementary table 1**: **Genetic map of the parental and Recombinant Inbred Lines used**. Matrix of the 101 lines used, 3 F1 heterozygotes (columns) and the 4,515 detected markers listed per chromosome (ch01-ch12) (lines). The genotypes are likelihood based where “0” indicates a locus derived from MM and “1” indicates a locus derived from PI. Chromosome number and genomic position (basepair) are given in the first two columns. Position is the average basepair position of the 100 SNPs sliding bin used to determine the parental origin of the locus.

**Supplementary table 2**: **Introgression size statistics**. Minimum, maximum, mean and median introgression sizes per chromosome (first column).

**Supplementary table 3**: Outcome of a linear model to detect differentially abundant mRNAs between the HP versus LN-treated RILs. For the mRNAs, two identifiers are given in the columns: identifier, and Name. Furthermore, the location (chromosome number and genome position of gene start in basepairs), orientation (+ or – strand), and size (length in basepairs) are indicated. Then, the outcome of the linear model is listed, first the significance in −log_10_(p) followed by two types of corrections for multiple testing: Bonferroni (conservative, as used in the main text) and Benjamini Hochberg False-discovery rate (FDR; less conservative for comparison). The column effect describes the difference between HP and LN treated maternal environment. A description of the effect direction (treatment) is given in the last column.

**Supplementary table 4**: Gene Ontology (GO) enrichment data of maternal environment-related mRNAs. Shown are the GO bin ID, GO bin category (GO name), GO aspect; molecular function (F), cellular component (C), and biological process (P), P value (p.value), Total mRNA’s identified in mRNA bin and GO (in.set), total number of genes in GO (In.GO) and total mRNA set size (set.size).

**Supplementary table 5**: Outcome of a linear model to detect differentially abundant mRNAs between the MM and PI parental lines and their interaction with the environment. For the mRNAs, two identifiers are given in the columns: identifier, and Name. Furthermore, the location (chromosome number and genome position of gene start in basepairs), orientation (+ or – strand), size (length in basepairs) are indicated. Then, the outcome of the linear model is listed, first the tested factor, then significance in −log_10_(p) and a correction for multiple testing (Benjamini Hochberg (FDR)). The column effect describes the difference between the factors tested and the interpretation of the effect direction is given in the last column.

**Supplementary table 6a**: Gene Ontology enrichment analysis of mRNAs that are higher in parental lines MM (left), PI (middle) and their interaction (right). Significantly different genes were taken from the model (see methods and material) only including the parental lines. See legend table S4 for details and abbreviations.

**Supplementary table 6b**: Gene Ontology enrichment analysis of mRNA differences between the parental lines and nutrient environment; higher in HP (left) or higher in LN (right). Significantly different genes were taken from the model (see methods and material) including the parental lines and the nutritional environment. See legend table S4 for details and abbreviations.

**Supplementary table 7a**: Heritability of mRNA abundances from the HP and LN maternal environments. The treatment column indicates the maternal nutrient environment, the mRNA ID is specified in the trait column. H2_keurentjes is the heritability, which was calculated using the genotypic variance (Vg) and the residual variance (Ve) as described in Keurentjes *et al*. (2007) [56]. The FDR column indicates the FDR = 0.05 threshold as determined by 1,000 permutations. The last two columns specify if an mRNA abundance was significantly heritable and whether it was specific for one or multiple maternal environments, or not (group).

**Supplementary table 7b**: Gene Ontology enrichment analysis of mRNAs with significant heritability.

**Supplementary table 8a**: Transgression for mRNA abundances from the HP and LN maternal environments in the RILs. The treatment column indicates the maternal environment, the mRNA ID is specified in the trait column. The n_lines_transgression column specifies how many RILs displayed transgression. The FDR column shows how many RILs showed transgression at the FDR = 0.05 threshold as determined by 1,000 permutations. The last two columns specify if an mRNA abundance was significantly transgressive or not and whether it was specific for one or multiple maternal environments, or not (group).

**Supplementary table 8b**: Gene Ontology enrichment analysis of mRNAs with significant transgression. See legend table S4 for details and abbreviations.

**Supplementary table 9**: Number of eQTLs detected per chromosome and treatment (LN; upper table, HP; lower table). Indicated are for all eQTLs, QTL type (*cis* or *trans*) and QTL effect found per chromosome per nutrient environment (+ or -). The last column indicates the number of eQTLs in the *trans*-bands (TB).

**Supplementary table 10**: List with eQTLs mapped in both the LN and HP maternal nutrient environments. First, the maternal environment is listed, second the mRNA ID (trait). Then columns with the location information of the eQTL: chromosome number, location (bp; and the confidence interval bp_left and bp_right), and the marker. Then, the significance in −log_10_(p) is given and the effect size (negative is higher in MM-derived loci; positive is higher in PI-derived loci). Furthermore, the type of QTL is given (*cis* or *trans*) and whether the QTL is part of a *trans*-band. Also, the variance explained by a single marker model is given (R2_sm). Subsequently, the name and location (chromosome number and start of the gene in basepairs) of the mRNA is listed.

**Supplementary table 11**: Gene Ontology enrichment in genes with an eQTL. See legend table S4 for details and abbreviations.

**Supplementary table 12**: Gene Ontology enrichment in genes with eQTLs mapping to a *trans-*band. See legend table S4 for details and abbreviations.

**Supplementary table 13**: Recombinant Inbred Lines per treatment.

**Supplementary figure 1**: Venn-diagrams showing **(A)** the overlap between all nutrient treatment-affected mRNA abundances and HP and LN heritable mRNA abundances and (**B**) mRNAs showing transgressive segregation in the HP and LN treatment.

**Supplementary figure 2**: Venn-diagrams showing the overlap between treatment-affected mRNAs, **(A)** *trans*-eQTLs and (**B)** *cis*-eQTL mapped in the HP and LN nutrient environments.

## Notes

### Competing Interest Statement

The authors have declared no competing interest.

