## Supplemental figures for "Plasticity of maternal environment dependent expression-QTLs of tomato seeds"

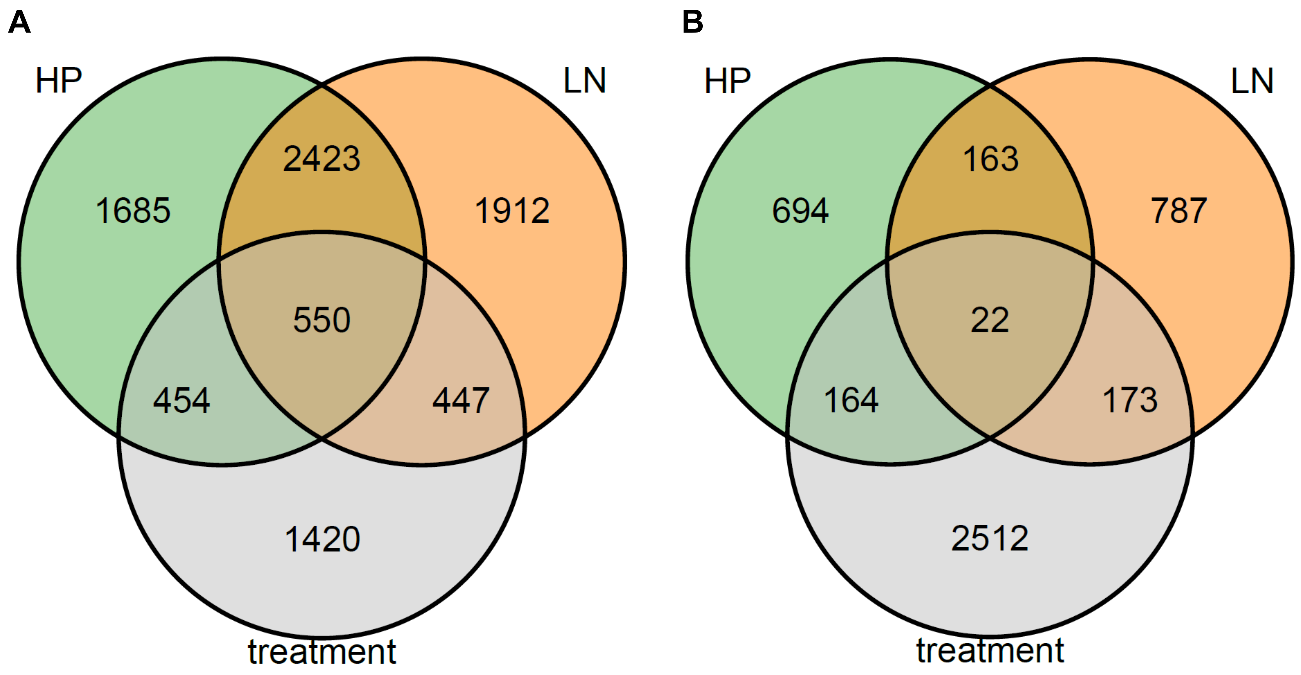


**Supplementary figure 1**: Venn-diagrams showing **(A)** the overlap between all nutrient treatment-affected mRNA abundances and HP and LN heritable mRNA abundances and (**B**) mRNAs showing transgressive segregation in the HP and LN treatment.

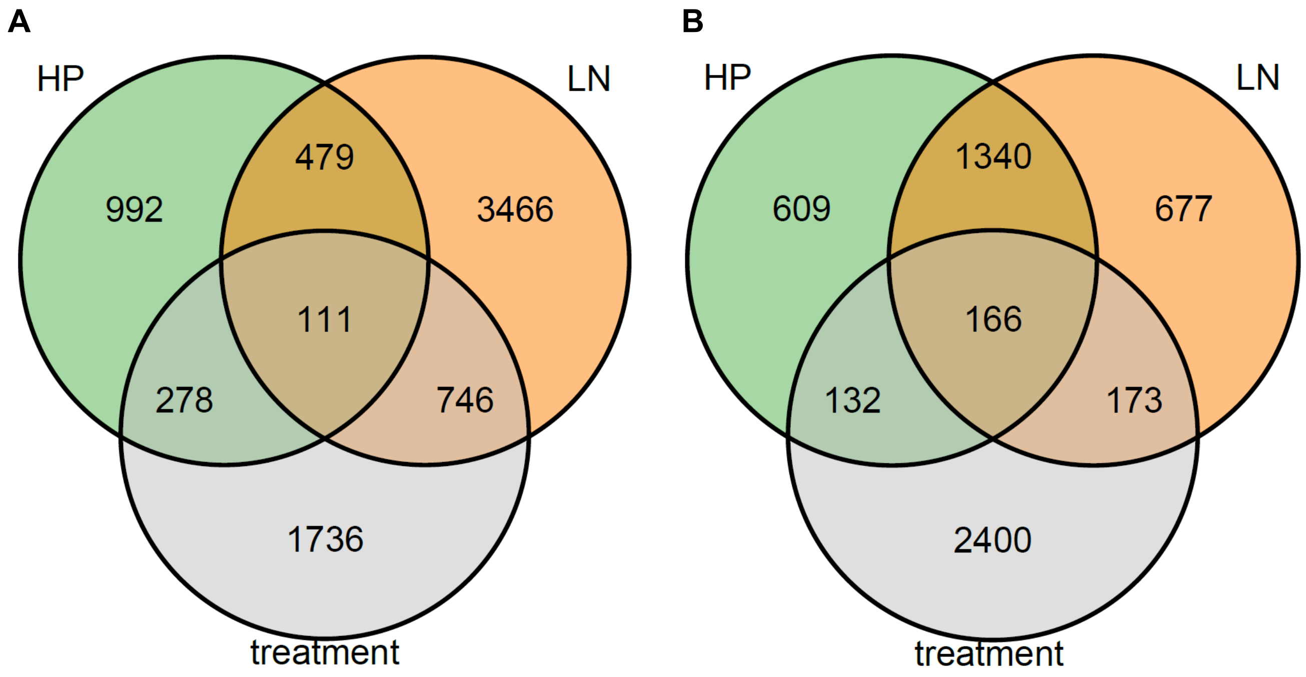


**Supplementary figure 2**: Venn-diagrams showing the overlap between treatment-affected mRNAs, **(A)** *trans*-eQTLs and **(B)** *cis*-eQTL mapped in the HP and LN nutrient environments.
